# The search for CDK4/6 inhibitor biomarkers has been hampered by inappropriate proliferation assays

**DOI:** 10.1101/2023.03.15.532719

**Authors:** Reece Foy, Kah Xin Lew, Adrian T Saurin

## Abstract

CDK4/6 inhibitors arrest the cell cycle in G1 and are used in combination with hormone therapy to treat advanced HR+/HER- breast cancer. To allow more effective use of these drugs in breast cancer, and to facilitate their use in other tumour types, biomarkers that can predict response are urgently needed. We demonstrate here that previous large-scale screens designed to identify the most sensitive tumour types and genotypes have misrepresented the responsive cell lines because of a reliance on ATP-based proliferation assays. When cells arrest in G1 following CDK4/6 inhibition, they continue to grow in size, producing more mitochondria and ATP. This cellular overgrowth masks an efficient arrest using metabolic ATP-based assays, but not if DNA-based assays are used instead. By comparing tumour cells using different assay types, we demonstrate that the lymphoma lines previously identified as the most responsive cell types, simply appear to respond the best because they fail to overgrow during the G1 arrest. Similarly, the CDK4/6 inhibitor abemaciclib appears to inhibit proliferation better than palbociclib, but this is because it also inhibits cell overgrowth through off-target effects. DepMap analysis of previous screening data using only the reliable assay types, demonstrates that palbociclib-sensitivity is associated with sensitivity to Cyclin D1, CDK4 and CDK6 knockout/knockdown, and resistance is associated with sensitivity to Cyclin E1, CDK2 and SKP2 knockout/knockdown. Furthermore, potential biomarkers of palbociclib-sensitivity are increased expression of Cyclin D1 (CCND1) and RB1, and reduced expression of Cyclin E1 (CCNE1) and CDKN2A. None of these associations are present when analysing DepMap using similar data from metabolic assays. This reinforces the importance of new screens to assess CDK4/6 inhibitors, and potentially other anti-cancer drugs, against a wide range of cell types using an appropriate proliferation assay. This would help to better inform clinical trials and to identify much needed biomarkers of response.

## INTRODUCTION

CDK4/6 Inhibitors are novel anti-cancer drugs that have revolutionised the treatment of breast cancer ^1, 2^. They arrest the cell cycle in G1 phase and are effective at treating advanced HR+/HER2- breast cancer, when used in combination with previous standard-of-care hormone therapy. CDK4/6 activity is required for G1 progression in many other cell types, implying that these drugs may also benefit a wider range of cancers. To identify the most sensitive tumour types, previous large-scale screens have assessed the effect of CDK4/6 inhibitors on the proliferation of a wide range of cancer cell lines ^3–6^. The aim of these screens is to reveal genomic features that correlate with sensitivity, thus yielding potential biomarkers of response. Predictive biomarkers are urgently needed, not just to define new tumour types that may be sensitive to CDK4/6 inhibitors, but to identify the breast cancer patients most likely to respond to these drugs ^7, 8^.

To date there have been numerous large-scale cancer line screens testing three licenced CDK4/6 inhibitors: palbociclib, abemaciclib and ribociclib. Palbociclib was included in the original Genomics of Drug Sensitivity in Cancer screen (GSDC1), which tested 403 compounds against 970 cancer lines ^3^. This reported that mutational inactivation of CDKN2A, which encodes for the endogenous CDK4/6 inhibitor p16^INK4A^, is associated with sensitivity to palbociclib. The subsequent GSDC2 screen assayed 297 compounds against 969 cancer lines, using a different ATP-based proliferation assay (CellTiter-Glo), and no association between CDKN2A loss/mutation and palbocicilib sensitivity was observed ^4^. Furthermore, the IC50 values were much higher overall in GDSC2, when compared to GDSC1 (cell types with sub-uM IC50: 21/968 in GDSC2 vs 123/888 in GDSC1).

The discrepancy between these screens is unclear, but a subsequent large-scale screen by Gong et al focussing exclusively on CDK4/6 inhibitors, produced similar results to GDSC2 using the same CellTiter-Glo endpoint ^5^. This screen tested palbociclib and abemaciclib against 560 cancer lines and could also not confirm the predictive value in CDKN2A loss/mutation. It did, however, identify other genomic aberration known to activate D-type cyclins as being associated with sensitivity to abemaciclib. Curiously, the IC50 values were also generally high for palbocicilb in the Gong et al study (22/560 lines with sub-uM IC50), although they were lower for abemaciclib (70/560 lines with sub-uM IC50). Furthermore, blood cancers were consistently the most responsive cancer types in both the Gong et al study and in GDSC2, and strangely, HR+/HER2- breast cancers appeared relatively insensitive ^4, 5^.

Finally, a molecular barcoding strategy known as PRISM, was recently used to characterise the response of 578 cancer lines to 4518 drugs ^6^. This large format was feasible because cancer lines containing DNA barcodes, which express unique mRNA transcripts, were screened together in pools. The pools were lysed 5 days after treatment and the relative abundance of each mRNA barcode was used to calculate the response of each cell type. This screen tested all three licenced CDK4/6 inhibitors, but none of these associated CDKN2A loss or mutation with sensitivity, nor identified any other potential biomarkers of response.

In summary, the GDSC1 screened appears somewhat of an outlier and most large-scale screens to date have failed to identify potential biomarkers of CDK4/6 inhibitor response. The exception is the Gong et al study, which identified genetic defects known to activate D-types cyclins, termed D-type cyclin activating features or DCAFs, which were predicted to be indicators of sensitivity to abemaciclib specifically ^5^. Unfortunately, these markers have yet to demonstrate predictive power in clinical studies.

We demonstrate here that a major problem with all of these screens is that they have used endpoints that do not directly measure proliferation. Instead, these endpoints measure the cumulative effects of cell number and cell size. This is particularly problematic for cells treated with CDK4/6 inhibitors, because these cells arrest in G1 but continue to grow in size ^9–13^. We show that this cell overgrowth causes scaling of mitochondria, thus cells appear to have “proliferated” using ATP-based endpoints, even though they have not. Cellular RNA similarly scales with growth, probably invalidating mRNA-based endpoints as well. We further demonstrate that the enhanced response observed in blood cancers or with abemaciclib, are due to reduce cell overgrowth under these conditions, and not due to an enhanced proliferative arrest. These misinterpretations have likely impeded the search for CDK4/6 biomarkers, because analysis of cumulative data from screens using only a reliable DNA-based assay, demonstrates expected markers of sensitivity and resistance. In particular CDKN2A loss is associated with sensitivity, whereas RB1 loss and Cyclin E overexpression are associated with resistance. This work calls for new screens to expand on this data using a reliable proliferation assay in a wide range of cancer cell lines. It also highlights the importance of using appropriate “proliferation” assays when assessing any anti-cancer drugs that arrests the cell cycle but permits continued cell growth.

## RESULTS

We recently demonstrated that CDK4/6 inhibition causes aberrant cell overgrowth during a G1 arrest, with cell size scaling linearly during a 4-day arrest period ^13^. This is broadly consistent with data from other groups, who also report increases in cell size following palbociclib treatment ^9–12, 14, 15^. We hypothesised that this cell growth could obscure the arrest using metabolic proliferation assays. To address this, we compared a panel of 8 cell lines using a range of different assays. Figure 1a demonstrates that 1μM palbociclib is sufficient to arrest most cell lines for up to 4 days, as assessed using live cell imaging to quantify cell cycle duration (mitosis to mitosis). This arrest is associated with cell growth throughout the arrest period in all lines (Figure 1b), as expected.

**Figure 1:**
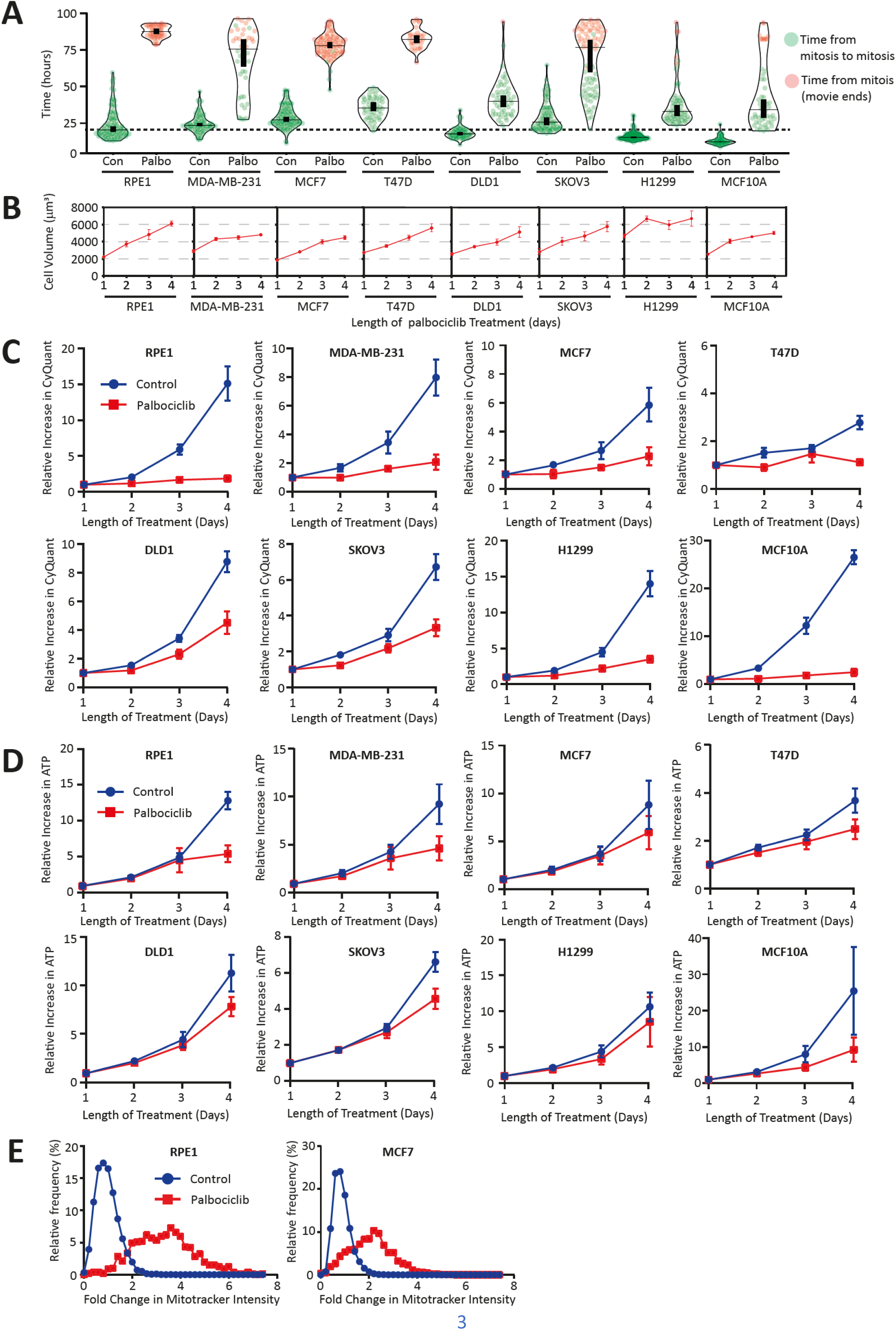
ATP assays fail to detect a proliferation arrest following CDK4/6 inhibition. **A)** Quantification of cell cycle length (mitosis to mitosis) of 8 different cell lines treated with DMSO (control) or palbociclib (1μM) and imaged continuously for 96hrs. Mitotic cells from the first 6hrs after treatment were selected and their daughter cells followed by eye until they reached mitosis or until the imaging period ended. Total cell cycle length was then recorded for each line in both the presence or absence of palbociclib. Horizontal lines show the median and thick verticals lines show 95% CI from 2 experiments, with at least 20 cells quantified per experiment. **B)** Cell volume assays of indicated cells following 1-4 days of palbociclib (1μM) treatment. Graphs display mean data +/- SEM from 3-4 repeats. **C)** CyQuant DNA quantification assays of indicated cell lines at 24hr intervals over a total of 4 days of treatment with DMSO (control) or palbociclib (1μM). Graphs display mean data +/- SEM from 3 repeats. **D)** Cell Titre Glo ATP quantification assays of indicated cells treated as in C. Graphs display mean data +/- SEM from 6 repeats. **E)** Frequency distribution of Mitotracker intensities from both RPE and MCF7 cells that were asynchronous (control) or treated with palbociclib (1μM) for 4 days. Graphs show combined data from 3 repeats, with at least 300 cells per condition.

We next compared the arrest using two commonly used proliferation assays that quantify either DNA content (CyQuant) or metabolic activity (CellTiter-Glo) as a surrogate for cell number. Figure 1C shows that CyQuant accurately detects a proliferation arrest in the presence of palbociclib, with data correlating well overall with the live-cell analysis (Figure 1A). In contrast, the metabolic CellTiter-Glo assay fails to detect the arrest in all cell lines, especially during the first 3 days of the assay which is the timepoint used in most large-scale screens (Figure 1D). We predicted that this was due to mitochondria scaling linearly with cell size during the arrest, thus increasing ATP output in the arrested overgrown cells. In agreement, mitochondria scaled with cell size in both of the two cell lines tested: RPE, and MCF7 (Figure 1E).

We next examined the effect of a dual PI3K and mTOR inhibitor, PF-05212384 ^16^, that we have previously shown can prevent cell overgrowth during a G1 arrest ^13^. PF-05212384 treatment combined with palbociclib inhibited mTOR activity and prevented G1 growth in all cell lines tested (Figure 2A,B). This combination did not markedly affect the proliferation assay using a DNA-based endpoints (Figure 2C), but it did improve the ability of the ATP-based assay to detect the arrest (Figure 2D). Our interpretation is that mTOR-mediated cell growth causes the production of new mitochondria in G1-arrested cells, which acts to obscure that G1 arrest using ATP-based assays. In support, inhibiting mTOR with PF-05212384 during a G1 arrest in either RPE or MCF7 cells also reduces the accumulation of mitochondria following palbociclib treatment (Figure 2E).

**Figure 2.**
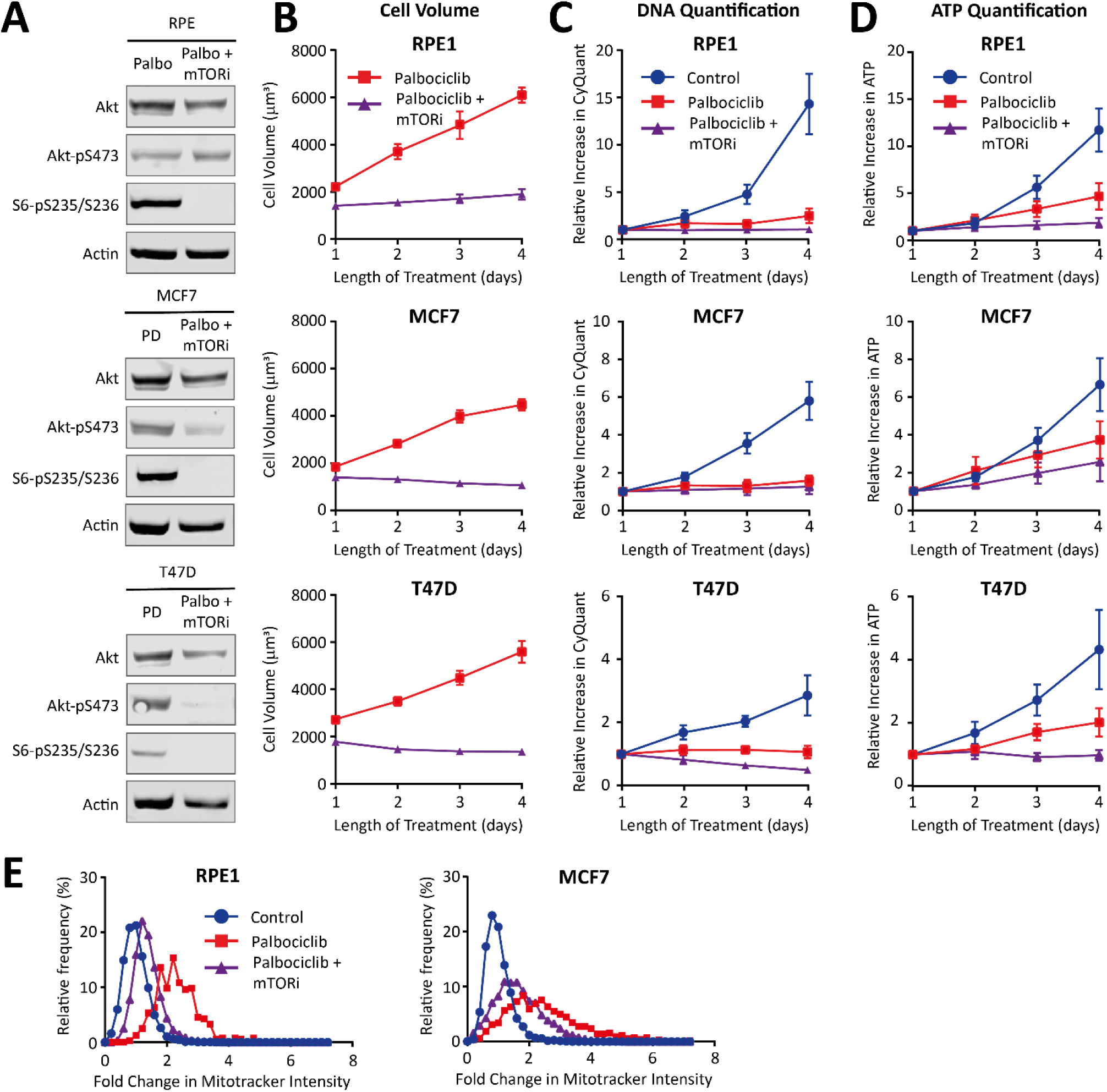
Cellular overgrowth obscures the CDK4/6 inhibitor arrest using ATP-based proliferation assays. **A)** Western blot analysis to analyse effect of the dual mTOR and PI3K inhibitor, PF-05212384, on mTOR activity using the downstream p70-S6K substrate S6-pS235/S236. Indicated cell lines were lysed following 24hrs of treatment with palbociclib (1μM) +/- PF-05212384 (30nM in RPE, and 7.5nM in MCF7/T47D). Blots are representative of 2 repeats. **B)** Cell Volume analysis in RPE, MCF7, and T47D cells following treatment with palbociclib (1μM) alone, or in combination with PF-05212384 (30nM RPE, 7.5nM MCF7/T47D) over 1-4 days. Graphs display mean data +/- SEM from 3-4 repeats. **C)** CyQuant DNA quantification of RPE, MCF7, and T47D cells treated as in B, or with DMSO (control). Graphs display mean data +/- SEM from 4 repeats. **D)** CellTiterGlo ATP quantification of RPE, MCF7, and T47D cells treated as in B and C. Graphs display mean data +/- SEM from 4 repeats. **E)** Frequency distribution of Mitotracker intensity from both RPE and MCF7 cells that were asynchronous (control) or treated with palbocilib (1μM) +/- PF-05212384 (30nM RPE, 7.5nM MCF7). Graphs show combined data from 2 repeats with at least 300 cells per condition.

Two previous large-scale screens have relied exclusively on the metabolic CellTiter-Glo assay: GDSC2 ^4^ and the Gong et all study ^5^, which remains the largest study to date profiling just CDK4/6 inhibitors. In contrast, the earlier GDSC1 screen used a DNA-based endpoint (Syto60) in a subset of adherent cells (645 cells total). We therefore compared these screens to search for differences that could reflect the different endpoints used.

Comparing IC50 values between GDSC1 and GDSC2 screens, demonstrates that sensitivity to palbociclib was considerably lower when using the metabolic CellTitreGlo endpoint (Figure 3A). Grouping tumours into their tissue of origin, demonstrates that this reduced sensitivity using metabolic assays holds true across all tumour types, consistent with data from Figure 1C and D. Interestingly however, blood cancers appeared to be considerably more sensitive to palbociclib using a metabolic assay. This was also observed for two different CDK4/6 inhibitors in the Gong et al study, which also used a metabolic endpoint (Figure S1). We therefore suspected this could reflect an unusual lack of cell growth in this tissue type. To test this, we randomly selected three lymphoma lines and tested their ability to arrest and grow in 1uM palbociclib. Figures 4A and B, demonstrates that all three lines arrested, based on cell counts, but failed to grow in size during that arrest. In support of a general inability of blood cancers to overgrow during a G1 arrest, a recent study similarly reported a lymphoma line as an outlier that failed to overgrow following palbociclib treatment ^15^. The lack of growth in the three lymphoma lines was associated with reduced mTOR activity during the G1 arrest (Figure 4C) and a lack of mitochrondrial scaling (Figure 4D). This was in stark contrast to MCF7 which actually increased mTOR activity (Figure 4C) and mitochondria (Figure 1E) during the arrest. The result is that the arrest was now detected efficiently in these lines using either a metabolic or DNA-based assays (Figures 4E,F).

**Figure 3.**
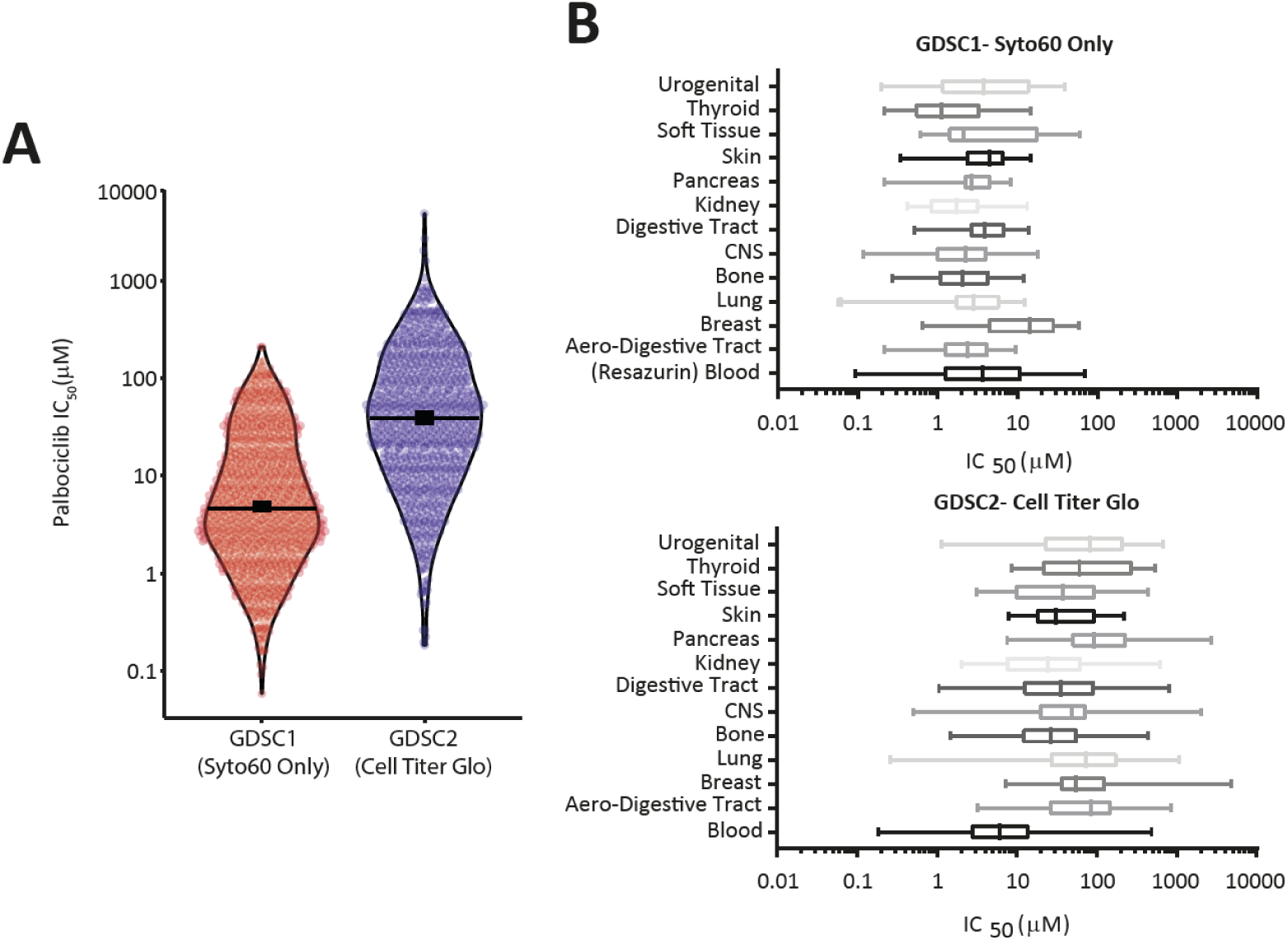
Previous large-scale screens using ATP assays report high IC50 values for palbociclib. **A)** Violin plot of all palbociclib IC50 values determined using Syto60 in GDSC1 against those measured using Cell Titer Glo in GDSC2. Horizontal lines show the median and thick vertical lines show 95% CI. **B)** Palbociclib IC50 values sorted into the tissue types from which the corresponding cancer line is derived. The top panel shows data from GDSC1 using the Syto60 DNA-based endpoint (except blood cancer shown with Resazurin), and the bottom panel shows all data from GDSC2 (CellTiter-Glo). The boxes represent the 25^th^ to 75^th^ percentile with the lines in the centre showing the median. Whiskers extend from the minimum to the maximum value for each dataset.

**Figure 4.**
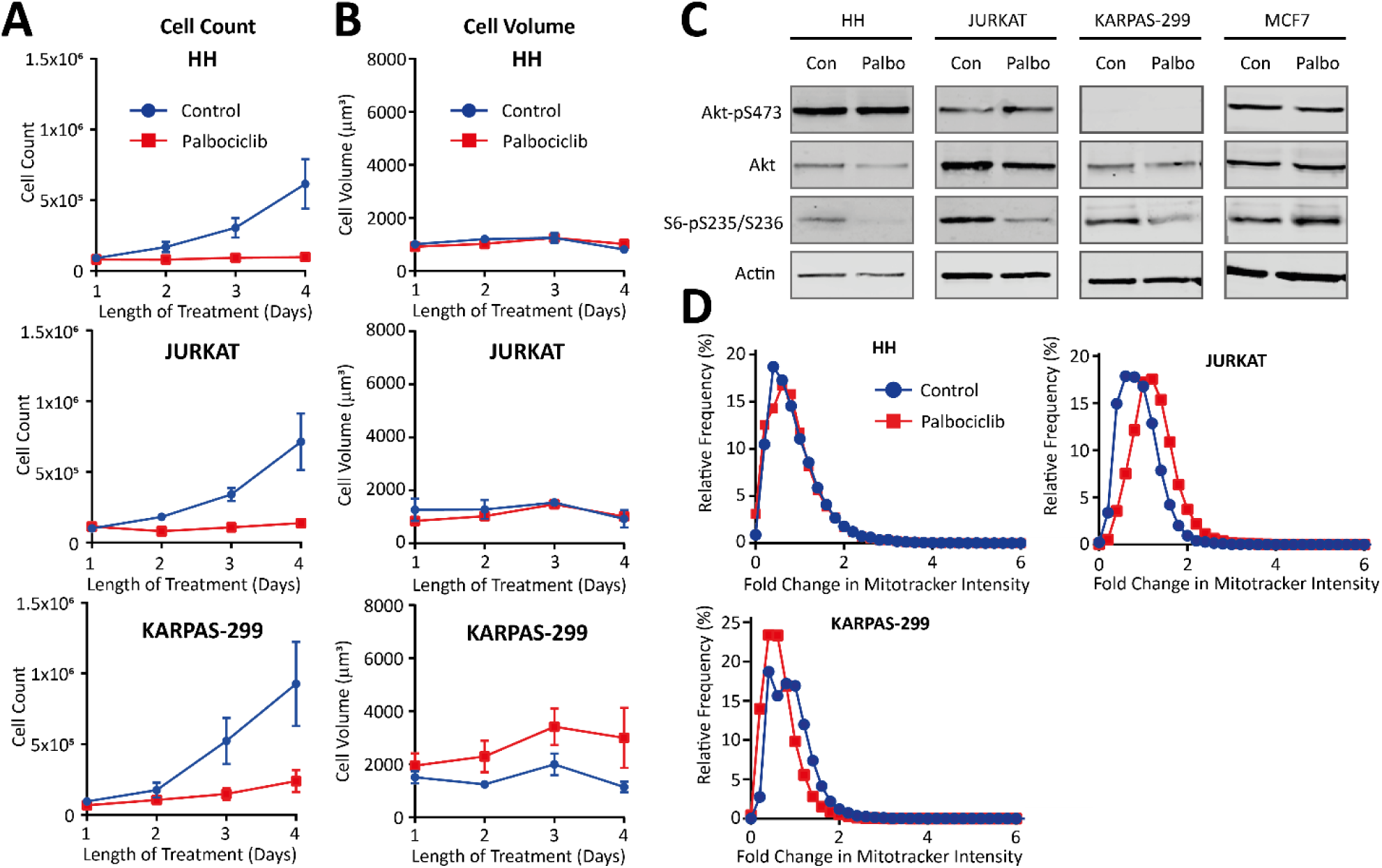
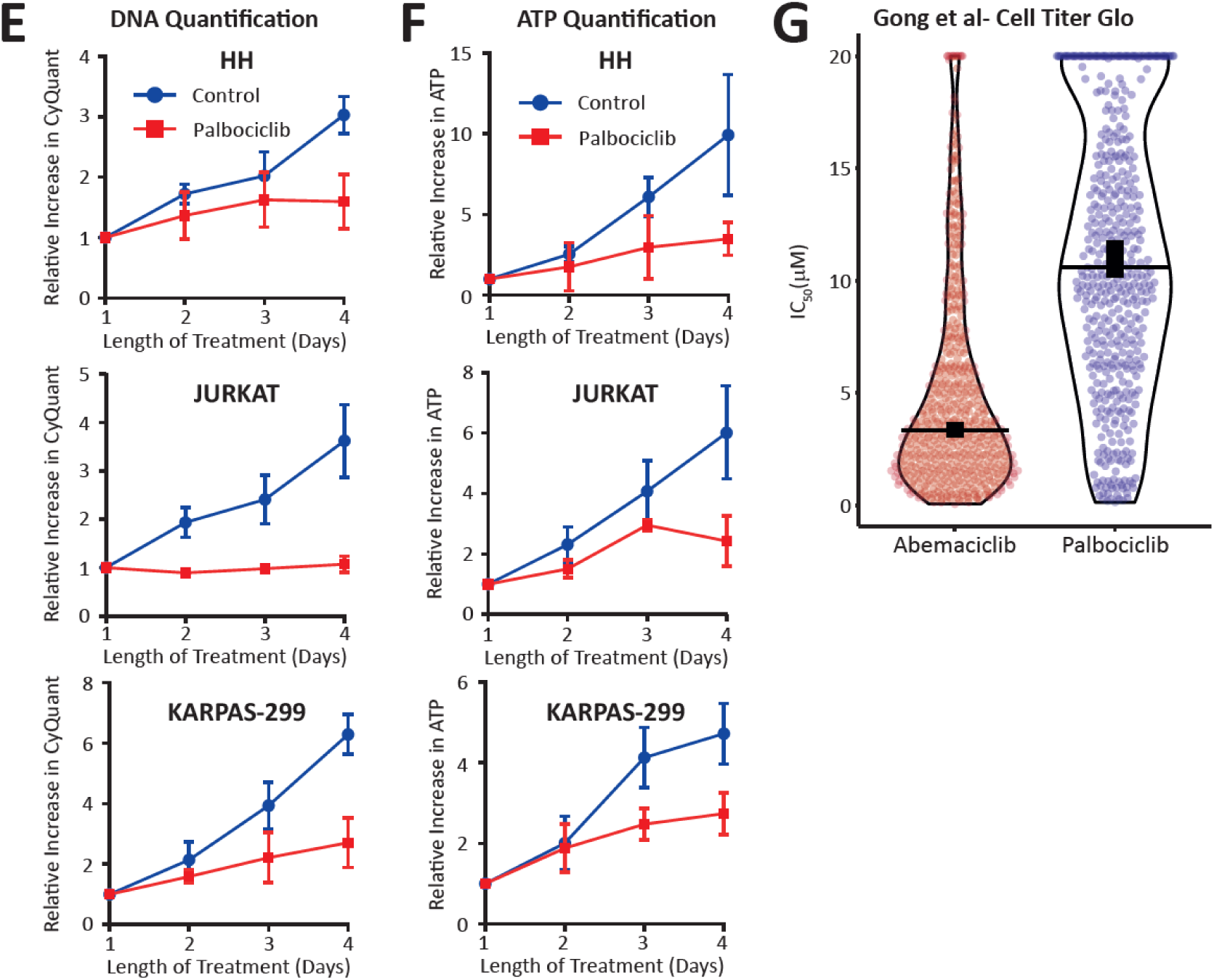
ATP assays accurately measure a proliferation arrest in lymphoma lines that fail to overgrow during the arrest. **A)** Quantification of HH, JURKAT, and KARPAS-299 cell number after treatment with DMSO (control) or palbociclib (1μM) for 1-4 days. Graphs display mean data +/- SEM from 4 repeats. **B)** Cell volume assays with the cells and treatments outlined in A. Graphs display mean data +/- SEM from 3-4 repeats. **C)** Western blot analysis showing mTOR activity in HH, JURKAT, and KARPAS-299 cells after 24hrs of treatment with DMSO (control) or palbociclib (1μM). Blots are representative of 2 repeats. **D)** Quantification of Mitotracker intensity in HH, JURKAT, and KARPAS-299 cells either untreated (control) or treated with palbociclib (1μM) for 4 days. Graphs contain data from 3 repeats, with at least 300 cells per condition. **E)** CyQuant DNA quantification of HH, JURKAT, and KARPAS-299 treated as in A and B. Graphs display mean data +/- SEM from 3-4 repeats. **F)** Cell Titre Glo ATP quantification of HH, JURKAT, and KARPAS-299 as in A, B, and E. Graphs display mean data +/- SEM from 3-4 repeats. **G)** Violin plot of IC50 values for either abemaciclib or palbociclib as determined by Gong et al using a CellTiterGlo ATP-based assay.

We also noticed that abemaciclib was generally more potent that palbociclib in the screen by Gong et al, which used a metabolic endpoint (Figure 4G) ^5^. Abemaciclib has also been shown to inhibit mTOR activity, through off-target effects on PIM1 ^17^. Therefore, we hypothesised that this compound could restrict growth during the G1 arrest. In agreement, abemaciclib-arrested cells displayed lower mTOR activity, reduced cell overgrowth, and restricted mitochondrial scaling in comparison to palbociclib-treated cells (Figure S2A-C). The net result is that abemaciclib appeared to arrest proliferation better than palbociclib specifically using an ATP-based assay, but not a DNA-based assay (Figure S2D-E).

In summary, ATP-based assays are unsuitable for measuring a G1 arrest following CDK4/6 inhibition because most cell types overgrow during that arrest and produce more mitochondria. The exception appears to be lymphoma lines, which shut off mTOR activity following CDK4/6 inhibition, thus preventing overgrowth and mitochondrial scaling. We suspect that these confounding effects of cell growth have also obscured data from the PRISM screens, which relied on barcodes encoded within mRNA transcripts ^6^. That is because total RNA also scales linearly during G1-overgrowth as a result of mTOR-mediated translation ^13^, and RNAseq demonstrates that most mRNA transcripts scale linearly with cell size/total RNA ^18^. Therefore, large, arrested cells, will also likely increase barcode levels and thus appear to have “proliferated” over the course of the assay despite remaining arrested in G1. This implies that the only partially reliable large-scale screen to date is GDSC1, which used a DNA-based endpoint in the adherent cell types, but a metabolic endpoint in the suspension cell types ^3^. In addition, a number of smaller scale screens have used either a DNA-based endpoints or cell counts to determine palbociclib sensitivity ^19–28^, which would both accurately determine arrest efficiency irrespective of cell size. We therefore combined data from these small screens with the GDSC1 adherent cell data and used DepMap to analyse co-dependencies (See Supplementary Table 1 for full list of cells lines and associated palbociclib IC50 data).

Figure 5A shows that the top 5 most significant co-dependencies using CRISPR or RNAi include Cyclin D1 (CCND1), CDK6, and CDK4, validating the palbociclib-sensitive cells as Cyclin D1-CDK4/6 dependent. Moreover, the top 3 most significant inverse co-dependencies for both CRISPR and RNAi are Cyclin E1 (CCNE1), CDK2 and SKP2, the E3 ligase that degrades p27 to activate CDK2 and promote S-phase. This demonstrates that resistant cells are preferentially reliant on Cyclin E1/CDK2 for S-phase entry, as expected ^29^. This analysis included a total of 686 cell lines, most of which were also assayed using a metabolic endpoint in large scale screens (567/686 lines). Analysing CRISPR or RNAi dependencies using the metabolic assay data does not extract these predicted co-dependencies (figure 5A), confirming that metabolic screens are unable to accurately detect CDK4/6 inhibitor sensitivity.

**Figure 5:**
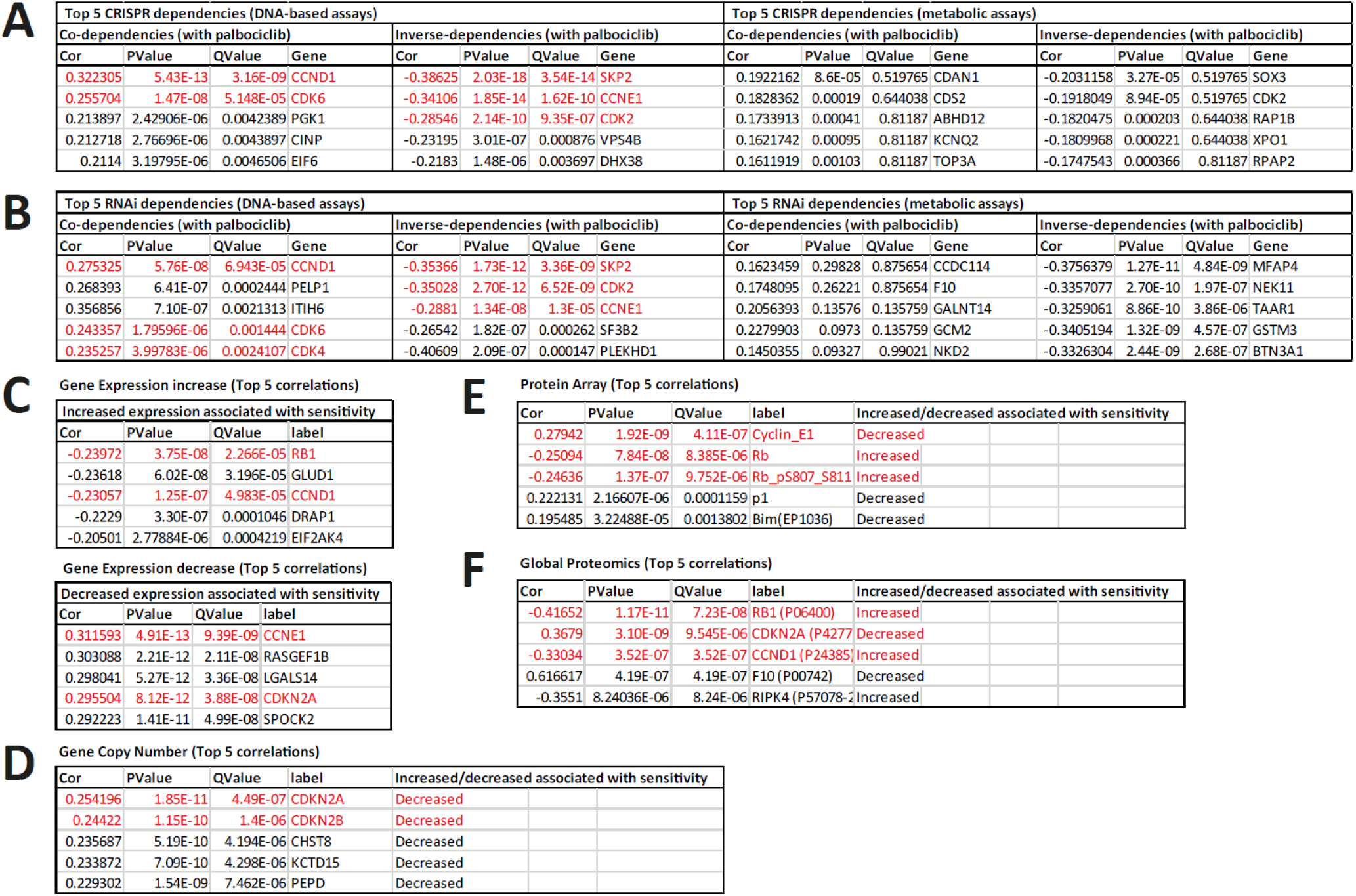
Reanalysis of previous screens using DNA-based endpoints reveals expected features of palbociclib sensitivity. **A-F)** DepMap Pearson correlation of palbociclib-sensitivity (using DNA based assays) against CRISPR dependencies (A), RNAi dependencies (B), Gene expression (C), Gene Copy Number (D), Protein Array (E) and Global Proteomics (F). Panels A and B also show DepMap Pearson correlation of palbociclib-sensitivity using metabolic assays for comparison. The top 5 most significant associations are shown for each group and the full top 1000 associations are shown in supplementary table 2.

Having validated the palbociclib-sensitivity data using DNA-based assays, we next sought to examine if this data could reveal potential biomarkers to help predict CDK4/6 response. Figure 5C shows that increased expression of RB1 and CCND1 is associated with sensitivity, whereas increased CCNE1 and CDKN2A is associated with resistance. Furthermore, similar changes are also detected as the strongest associations with gene copy number and total protein levels (Figure 5D-F). The full DepMap correlation data is display in supplementary table 2. In summary, this analysis of previous data, generated using reliable DNA-based proliferation assays, extracts many of the genomic features predicted to lead to CDK4/6 sensitivity or resistance. The also agree with the earlier findings that CDKN2A loss or mutation are potential biomarkers of sensitivity ^3, 21^.

## DISCUSSION

We demonstrate here that most cells continue to grow in size when they are arrested by CDK4/6 inhibitors. These enlarged cells scale their mitochondria and therefore appear to still be proliferating using ATP-based assays, even though they are not. We have shown recently that total cellular protein and RNA also scales with size during the arrest ^13^, therefore we hypothesise that “proliferation” assays that use any of these endpoints will also misrepresent cell enlargement as cell proliferation. This affects every large-scale screen that has been carried out to date to assess CDK4/6 inhibitor sensitivity. The GDSC1 screen relied partially on ATP-assays ^3^, and the GDSC2 and Gong et all screens relied exclusively on ATP-assays ^4, 5^. The pooled PRISM assays, which screened all three licenced CDK4/6 inhibitors, relied on mRNA sequencing of lentiviral barcodes to determine the relative levels of each cell line within the pools before and after treatment ^6^. It is likely that these barcodes also scale with size, as do most cellular mRNAs during a CDK4/6 inhibitor arrest ^13, 18^. Therefore, we predict that overgrown cells will similarly be overrepresented in PRISM assays despite an effective arrest. There is therefore an urgent need to perform new assays that can accurately report a proliferative arrest following CDK4/6 inhibitor treatment.

The same issue may affect the assessment of many more anti-cancer drugs. That is because an arrest at different cell cycle stages also causes cells to overgrow. This was shown recently for the CDK7 inhibitor ICEC0942 ^30^, which arrests cells in G1 and G2, and we have observed continued growth after a thymidine-induced S-phase arrest (data not shown). Therefore, anti-cancer drugs that indirectly halt cell cycle progression by causing DNA damage are likely to similarly cause overgrowth in the arrested cells. Finally, cell enlargement is a hallmark of senescence ^31–33^, therefore if cells permanently exit the cell cycle following drug treatment, these cells will similarly be overrepresented using most proliferation assays. Together, this could explain why the cellular IC50 values for a wide range of anti-cancer drugs were previously shown be artificially high using metabolic assays, in comparison to a DNA based endpoint ^34^. This reinforces the need to use proliferation assays that are not influenced by cell size when assessing anti-cancer drugs.

We propose that an ideal assay would independently measure both cell cycle arrest and cell size. That is because cell overgrowth drives toxicity and cell cycle exit following CDK4/6 inhibition ^13^, an affect that is also seen following CDK7 inhibition ^30^. Therefore, cells that arrest efficiently and overgrow the most may ultimately respond the best to cell cycle inhibitors. The mechanistic explanation is that overgrown cells experience osmotic stress during G1, causing p21 induction and a fraction of cells to enter senescence ^14^. Cells that escape this arrest enter S-phase and experience further DNA damage as a result of replication stress ^18^ and a weakened DNA damage checkpoint ^15^. This causes further cell cycle arrest from G2 or catastrophic DNA damage during mitosis as under-replicated chromosomes are segregated. It will be important in future, to determine if these toxic effects of cell overgrowth after a cell cycle arrest, cause senescence following treatment with a wide variety of anti-cancer drugs. This could explain why cell enlargement is a hallmark of senescence ^31–33^, and that would further reinforce the need to accurately assess this enlargement when treating with drugs that are predicted to induce senescence.

It is unclear why blood cancers fail to grow during a palbociclib arrest, but the prediction is that these cancer types will be the least sensitive to downstream toxicity and cell cycle withdrawal. The fact that these cells have been categorised as some of the most sensitive by current assays, further underscores the importance of using the correct endpoint in these assays. Similarly, abemaciclib has been proposed to inhibit proliferation better than palbociclib, when in fact, this is due to its ability to limit overgrowth through off-target effects (Figure S2). Whether this is beneficial in the treatment of HR+/HER2- breast cancer is currently unclear, but in cell models at least, it likely explains the rather limited cell cycle withdrawals following abemaciclib treatment, in comparison to three other licenced CDK4/6 inhibitors ^18^.

All of these confounding effects of cell growth have likely hampered the search for CDK4/6 biomarkers. By grouping and analysing assays using reliable endpoints, we find that the top co-dependencies are Cyclin D1, CDK4 and CDK6 for palbociclib-sensitive cells, and Cyclin E1, CDK2 and Skp2 for palbociclib-resistant cells (Figure 5a). Comparisons of gene expression, copy number and proteins levels identified decreased CDKN2A as one of the strongest a predictors of sensitivity (Figure 5D, E). This gene encodes for the CDK4/6 inhibitor p16 ^INK4A^, and low p16 ^INK4A^ expression was previously identified as a possible marker of sensitivity to palbociclib ^21^. The subsequent PALOMA-1 trial showed CDKN2A copy number was not predictive of response ^35^, and the later PALOMA-2 trial showed CDKN2A mRNA or expression of p16^INK4A^ protein were also not predictive either ^36^. However, as discussed here ^37^, the ER+/HER2- patients in these trials may already have low CDKN2A levels, thus removing its predictive power. Furthermore, high CDKN2A was recently shown to be a biomarker of resistance to CDK4/6 inhibitors in ER+ breast cancer ^38^. A rationale for how CDK4/6 inhibitor proteins can drive resistance was recently provided by the demonstration that p16^INK4B^ and p18^INK4C^ preferentially associate with CDK6 and distort the ATP binding pocket such that it favours ATP binding over palbociclib ^39^. Therefore, inhibitory INK4 proteins may need to be kept low if CDK6 is expressed to ensure sensitivity to CDK4/6 inhibitors. Whether CDKN2A or CDKN2B loss drives sensitivity specifically in cell types that overexpress CDK6, remains to be determined. These and other crucial biomarker questions will ultimately be facilitated by new screens that assess CDK4/6 inhibitor sensitivity in a large number of cancer cell lines using a reliable DNA-based assay type.

## ACKNOWLEDGEMENTS

This work was funded by a Tenuvus PhD studentship to RF, and a Cancer Research UK Programme Foundation Award to ATS (C47320/A21229), which also funds RF. The authors declare no competing financial interests.

## MATERIALS AND METHODS

### Cell Culture and Reagents

hTERT-RPE1 (RPE1), MCF7, T47D, H1299, MCF10A, MDA-MB-231, Jurkat (clone E6-1), and HH cells were from ATCC. The human ovarian adenocarcinoma SKOV3 was acquired from CRUK. The human colorectal cancer line DLD1-FRT was a kind gift from Stephen Taylor, which was published previously ^40^. The human non-Hodgkins Ki-positive large cell lymphoma KARPAS-299 was from public health England. All cells were validated by STR profiling and periodically checked to confirm they were mycoplasma free.

All cells were cultured at 37°C with 5% CO2. RPE1, MCF7, T47D, DLD1-FRT, SKOV3, and MDA-MB-231 were cultured in DMEM (Thermo Fisher Scientific, Gibco 41966029) supplemented with 9% FBS (Thermo Fisher Scientific, Gibco 10270106) and 50 μg/ml penicillin/streptomycin. MCF10A cells were cultured in F12/DMEM (Thermo Fischer Scientific, Gibco, 11320033) and supplemented with 5% horse serum (Thermo Fischer Scientific, Gibco 16050122), 20ng/mg EGF (Sigma, E9644), 0.5ug/ml hydrocortisone (Sigma, H088), 100ng/ml cholera toxin (Sigma, C8052), 10ug/ml insulin (Sigma, I9278) and 50ug/ml penicillin/streptomycin (Sigma, P4458). H1299, HH, Jurkat, and KARPAS-299 were cultured in RPMI-1640 (Sigma Life Science, R8758) supplemented with 9% FBS and 50 μg/ml penicillin/streptomycin. Palbociclib (PD-0332991) was purchased from MedChemExpress (HY-50767A) and PF-05212384 (Gedatolisib) was purchased from Sigma (PZ0281). Abemaciclib (LY-2835219) was purchased from Selleckchem (S7440).

### Time Lapse Imaging

To characterise the arrest caused by palbociclib each cell line was plated at low density (15,000 per well) into an Ibidi μ-plate glass-bottomed 24 well plate. The following day cells were treated with drugs and then imaged using a Holomonitor M4 (Phase Holographic Imaging) at 37°C with 5% CO2. Images were taken every 20 minutes for a total of 4 days. Image analysis was performed using the Holomonitor App Suite. For each condition, cells were selected at random and then followed by eye to record the length of time between the first and second mitosis (or the end of the movie).

### Cell Volume Analysis

For cell volume analysis, cells were plated into 6-well plates at a density of 30,000 cells per well (RPE, MCF7, T47D, MDA-MB-231, DLD1, SKOV3, H1299, MCF10A) or 100,000 cells per well (HH, JURKAT, KARPAS-299) then treated with drugs immediately, and incubated at 37°C with 5% CO2. At 24hr intervals, cells were trypsinised, stained with acridine orange and DAPI, and cell diameters were measured using a Chemometec NC-3000 Nucleocounter. Histograms of cell diameter were imported into Flowing Software version 5.2.1 and mean cell diameter of each condition was taken to calculate volume using 4/3 πr^3^.

### Cell Proliferation Assays

Cell lines were plated at a density of 1000 cells per well (RPE, MCF7, T47D, MDA-MB-231, SKOV3, DLD1, H1299, and MCF10A) or 20,000 cells per well (HH, JURKAT, KARPAS-299) in a 96-well plate and immediately treated with drugs and incubated at 37°C with 5% CO2. Cell proliferation was then measured by different means at 24hr intervals for a total of 4 days. DNA was quantified using CyQuant Direct Cell Proliferation assay (ThermoFischer Scientific, C35011). A 2x CyQuant solution (1/50 background suppressor and 1/250 nucleic acid in phosphate buffered saline) was added directly to the cells and incubated at 37°C with 5% CO2 for 30 minutes. The plate was then removed from incubation and fluorescence intensity was read at 485/535nm using a Tecan Infinite F Plex plate reader. ATP was then quantified using CellTiter-Glo Luminescent Cell Viability Assay (Promega), either in the same cells that were stained with CyQuant or in a set of separate but identical samples. Prior to ATP quantification all cells were briefly washed in PBS then a solution containing equal parts CellTiter-Glo reagent and growth medium was added to each well and incubated at room temperature for 15 minutes. Samples were then transferred to a white-bottomed 96-well plate, and luminescence was read using a Tecan Infinite F Plex plate reader. For direct cell counts, HH, JURKAT, and KARPAS-299 were plated at a density of 100,000 cells per well, treated with drugs immediately, and incubated at 37°C with 5% CO2. At 24hr intervals, cells were harvested, stained with acridine orange and DAPI, and then counted using a Chemometec Nucleocounter NC-3000.

### Mitochondrial quantifications

Cells were plated at 30,000 (RPE, MCF7) or 100,000 (HH, JURKAT, KARPAS-299) cells per well in a 6-well plate, immediately treated with drugs, and incubated at 37°C with 5% CO2 for 4 days. Total mitochondrial mass per cell was then quantified using MitoTracker Deep Red FM (ThermoFischer Scientific, M22426). Each well was treated with 500nM of mitotracker reagent and incubated at 37°C with 5% CO2 for 30 minutes. Cells were then trypsined, stained with acridine orange, and fluorescence was measured with a Chemometec NC-3000 Nuceocounter using a Flexicyte analysis algorithm. Cell masking was achieved using darkfield microscopy, and acridine orange staining (530nm) was used as counterstaining to ensure that only the DNA positive events identified in the darkfield were analysed for MitoTracker staining (630nm).

### Western Blotting

Protein lysates for western blotting were prepared by scraping cells into 4x sample buffer (250mM Tris, 10% SDS, 40% Glycerol, 0.1% Bromophenol Blue). Lysates were then sonicated (15 second pulse 50% amp) with a Cole-Palmer Ultrasonic Processor. Samples were then briefly boiled and centrifuged at max rpm for several seconds. Protein concentration was determined via DC assay, after which samples were diluted to desired concentration in sample buffer and 2-mercaptoethanol was added to a final concentration of 10%. Equal volumes of sample were then separated by SDS-PAGE and transferred to 0.45um nitrocellulose membranes (Amersham Protran Premium). Membranes were then blocked for 15 minutes with 5% BSA in TBS with 0.1% Tween 20 (TBS-T) then incubated at 4°C overnight in 5% BSA TBS-T containing primary antibodies. The following day membranes were washed three times with TBS-T before being transferred to secondary antibodies in a solution of 5% milk in TBS-T. After 2 hours of incubation at room temperature membranes were washed three more times in TBS-T and imaged on a LI-COR Odyssey CLx system. The following primary antibodies were used for western blotting: rabbit anti-AKT (Cell Signaling Technology, 9272, 1/1000), rabbit anti-pAKT (Ser473)(Cell Signaling Technology, 4060, 1/1000), rabbit-pS6 (Ser235/236)(Cell Signaling Technology, 4856, 1/1000), and rabbit anti-actin (Sigma, A2066, 1/5000). The secondary antibody used was IRDye 800CW Goat anti-Rabbit IgG (LI-COR, 1/15000).

### DepMap analysis

A literature search identified 11 studies that used DNA-based or cell count-based proliferation assay to determine palbociclib IC50s ^3, 19–28^. The data from these studies was combined, and duplicate cell lines data was removed. The criteria for removing duplicates was that if the same line was screened by multiple studies the smaller scale screening data was used in preference (i.e. in preference to the large-scale GDSC1 data). If duplicate data from more than 1 small scale screens was present, we retained the data that recorded the lowest IC50. A Pearson correlation was then computed using Depmap (http://depmap.org) against CRISPR dependencies, RNAi dependencies, gene expression, gene copy number, protein array and proteomics. The full list of correlations, with associated P values and Q values, is displayed in Supplementary table 2, and the top 5 most significant associations is displayed in Figure 5. To compare how analysis of the same cell lines performed using metabolic assays, we extracted data from the GDSC2 screen ^4^, which used a metabolic endpoint and included most of the cell lines screened using ATP assays. For any that were not included in GDSC2, metabolic data was extracted from the Gong et al screen ^5^ or the GDSC1 suspension cell data ^3^. Out of a total of 686 lines screened with a DNA endpoint, 567 were also screened with metabolic endpoints. The full metabolic screening data is included in supplementary table 3. Using this data, a Pearson correlation was computed using Depmap against CRISPR dependencies and RNAi dependencies, and the full list of correlations is displayed in supplementary table 2, with the top 5 most significant associations displayed in Figure 5B.

### Statistical analysis

PlotsofData was used to make violin plots at https://huygens.science.uva.nl/PlotsOfData/ ^41^. These plots display the 95% confidence intervals (thick vertical bars) calculated around the median (thin horizontal lines). This allows statistical comparison between all conditions on the plots because when the vertical bar of one condition does not overlap with one in another condition the difference between the medians is considered statistically significant (p<0.05).

## Notes

### Competing Interest Statement

The authors have declared no competing interest.

### Summary of Updates

small corrections to text

