## Supplementary Figures for "The search for CDK4/6 inhibitor biomarkers has been hampered by inappropriate proliferation assays"

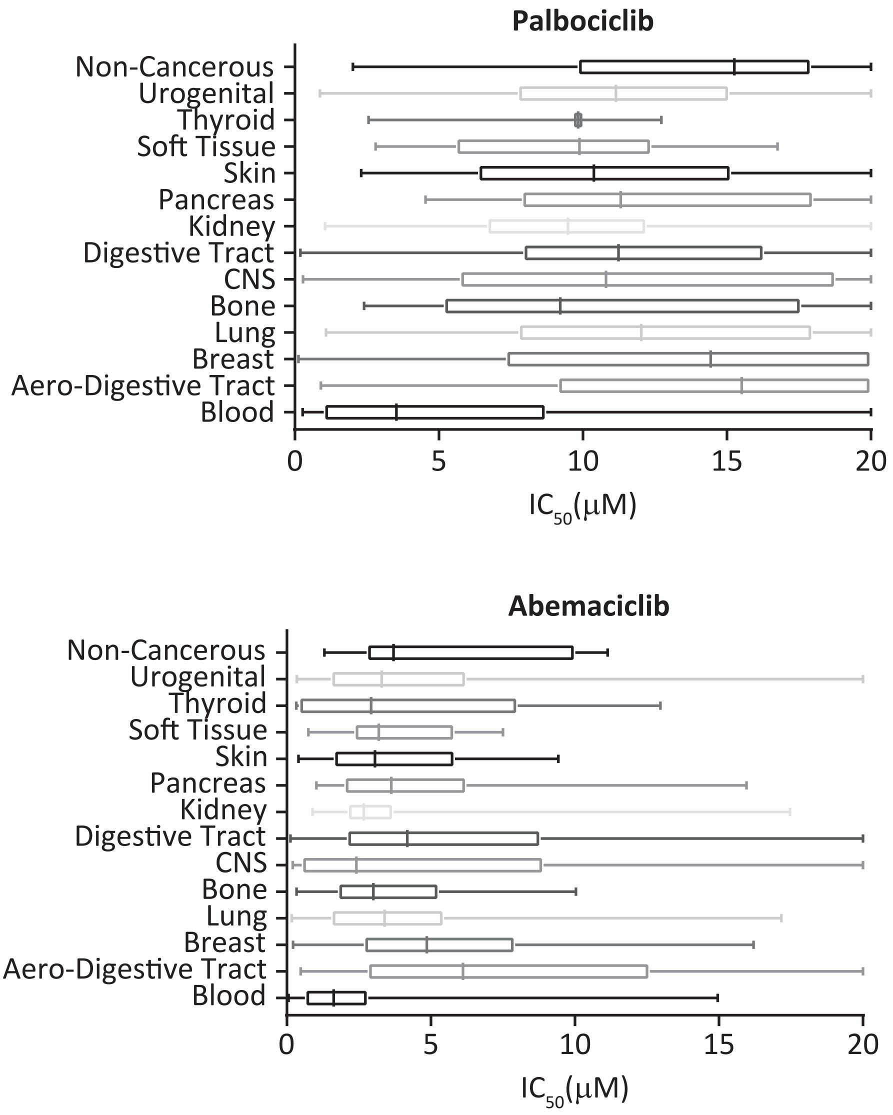


**Figure S1- A previous metabolic screen found blood cancers to be uniquely sensitive to CDK4/6 inhibitors.** IC50 values for either palbociclib or abemaciclib as determined in Gong et al 2017 using CellTiter-Glo ATP assays. Values are sorted into tissue types from which individual cancer lines are derived. The boxes represent the 25th to 75th percentile with the lines in the centre showing the median. Whiskers extend from the minimum to the maximum value for each dataset.

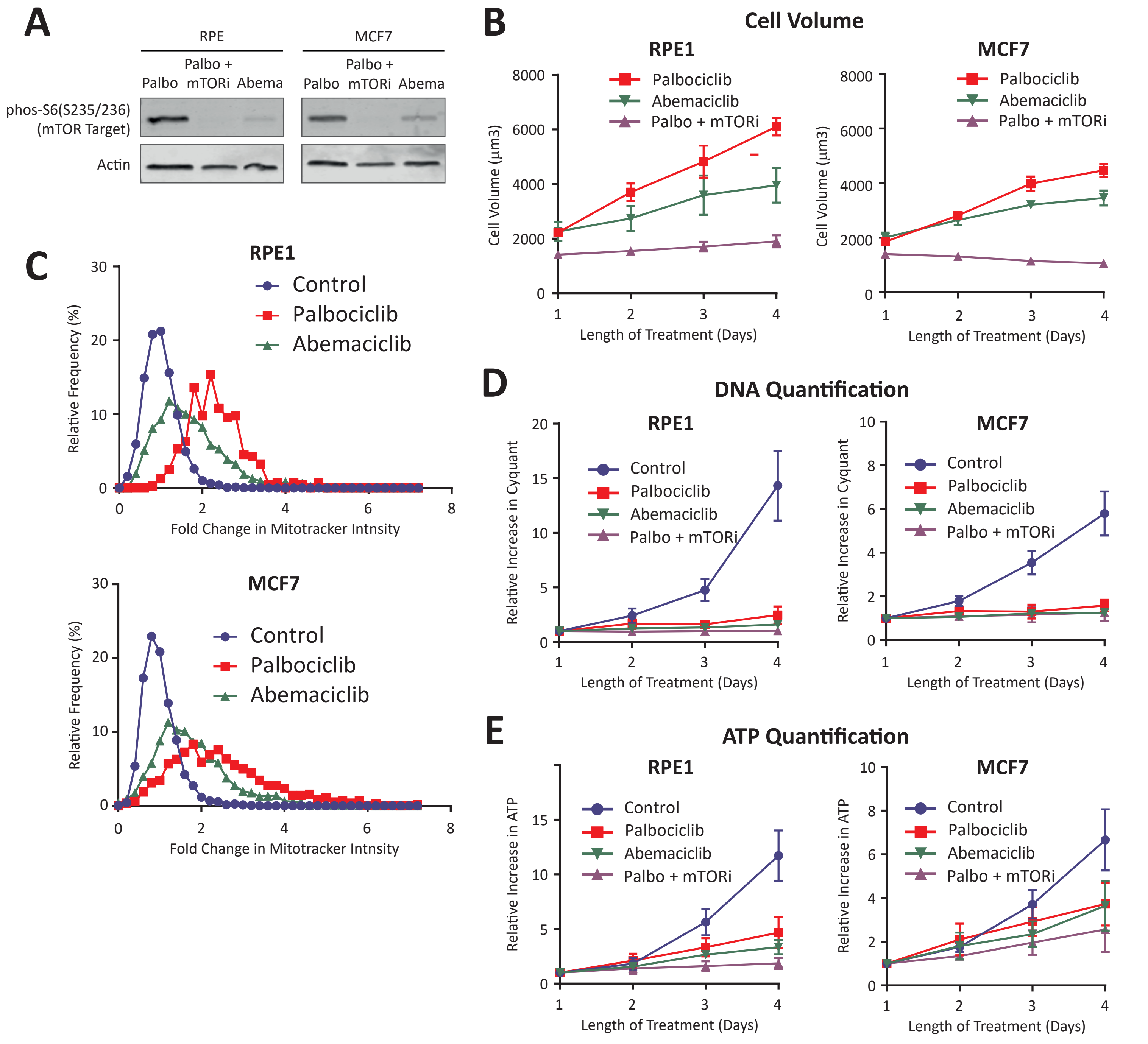


**Figure S2- Abemaciclib partially inhibits mTOR, thus restricting overgrowth and mitochondria scaling.**
**A.** Western blot analysis showing mTOR activity in RPE1 and MCF7 cells treated with palbociclib (1µM) +/- PF-05212384 (30nM in RPE, and 7.5nM in MCF7), or abemaciclib (600nM) for 24hrs. Blots are representative of 2 repeats.
**B.** Cell volume analysis of RPE1 and MCF7 cells treated with abemaciclib (600nM) for 1-4 days. Error bars represent +/- SEM from 3 repeats. Data for palbociclib (+/- PF-05212384) has been taken from figure 2B for comparison.
**C.** Quantification of mitotracker intensity in either RPE1 or MCF7 cells treated with abemaciclib (600nM) for 4 days. Graphs display data from 2 repeats. Control and palbociclib data were taken from figure 2E for comparison.
**D.** Quantification of DNA (CyQuant) and ATP (CellTiter-Glo) in RPE1 and MCF7 cells treated with abemaciclib (600nM) for 1-4 days. Error bars represent +/- SEM from 3 repeats. Control and palbociclib data were taken from figures 2C and 2D for comparison.
